# Rational design of immune gene therapy combinations via *in vivo* CRISPR activation screen of tumor microenvironment modulators

**DOI:** 10.1101/2023.03.14.532665

**Authors:** Guangchuan Wang, Feifei Zhang, Ryan D. Chow, Emily He, Lvyun Zhu, Qin Han, Sidi Chen

## Abstract

The hostile tumor microenvironment (TME) is major challenge for cancer immunotherapies. Here, we design and perform TME-targeted *in vivo* CRISPR activation (CRISPRa) screens to uncover factors that promote anti-tumor immunity, culminating in rationally designed immune gene therapy combinations. Through adeno-associated virus (AAV) delivery, multiplexed activation of pooled immunoregulatory genes encoding <u>a</u>ntigen <u>p</u>resentation, <u>c</u>ytokine, and co-stimulation <u>m</u>olecules (APCM) leads to enhanced anti-tumor immunity. APCM screen in metastatic tumors identifies *Cd80, Tnfsf14, Cxcl10, Tnfsf18, Tnfsf9*, and *Ifng* as the top immunostimulatory candidates. AAV-mediated delivery of these factors individually or in combination shows anti-tumor efficacy across different cancer models. Further optimization pinpoints *Ifng*+*Tnfsf9*+*Il12b(Il12/Il23)* as a potent therapeutic combination, leading to increased IFN-γ^+^CD8^+^ and tissue-resident memory T cells. APCM therapy synergizes with CAR-T cell therapy against human solid tumors *in vivo*. APCM-based CRISPRa screen and gene activation systems can thus be leveraged for the rapid generation of off-the-shelf immune gene therapies against solid tumors.

## Introduction

Immune checkpoint blockade (ICB) (Pardoll, 2012; Sharma and Allison, 2015) and adoptive cell therapies (June et al., 2018a; Kershaw et al., 2013; Stadtmauer et al., 2020) have successfully turned some late-stage, metastatic, and chemo-resistant lethal cancers into manageable diseases in a subset of patients (Ribas and Wolchok, 2018; Rosenberg and Restifo, 2015; Sharma and Allison, 2015; Sharma et al., 2017). The clinical success of ICB has transformed cancer therapeutics (Chen and Mellman, 2013; Mellman et al., 2011). Nevertheless, the majority of the patients still do not benefit from these immunotherapies (Chen and Mellman, 2017; Sharma et al., 2017). Adoptive cell therapies with genetically engineered chimeric antigen receptor (CAR, referred as CAR-T cells), have demonstrated clinical success in the treatment of multiple types of liquid cancers such as leukemia, lymphoma and multiple myeloma (June et al., 2018b; Ghorashian et al., 2019; Rafiq et al., 2019; Raje et al., 2019). Extensive efforts have been devoted into the development of T cell therapies for various solid tumors, including *in vitro* expanded tumor-infiltrating lymphocytes (TILs), CAR-T cells, and genetically engineered T cell receptor (TCR) T cells (Liu et al., 2021; Ma et al., 2019, 2020; Morotti et al., 2021; Vodnala et al., 2019). However, adoptive T cell therapies with TILs, CAR-T cells or TCR-T cells have not consistently achieved clinical success against solid tumors. The failure of cell-based immunotherapies to date against solid tumors demonstrates that an abundance of tumor-reactive circulating T cells (exogenously introduced or not) is insufficient for sustained immune responses against solid tumors. Corroborating the results of cell-based therapies, studies employing therapeutic cancer vaccines have demonstrated that the generation of tumor-reactive immune cells is only one piece of the larger puzzle (Joyce and Fearon, 2015; Rosenberg et al., 2005). In particular, while both traditional cancer vaccines and personalized peptide-or RNA-vaccines targeting neoantigens can induce large numbers of tumor-specific T cells in the periphery, these vaccines generally fail to elicit robust anti-tumor effects in isolation (Palucka and Banchereau, 2012; Rosenberg et al., 2004; Sahin and Türeci, 2018; Saxena et al., 2021).

A fundamental difference between solid and liquid cancers is the specialized, immune-privileged tumor microenvironment (TME) (Galon and Bruni, 2019; Joyce and Fearon, 2015; Kaymak et al., 2020). The success of anti-tumor immunity requires not only high frequencies of cancer-specific T cells that can be addressed by adoptive cell therapy and cancer vaccine, but also the effective infiltration and maintenance of functional immune cells within TME. Overcoming the immune-privileged TME is thus a primary focus in the development of next-generation immunotherapies, potentially by modulating the TME (Chapman et al., 2020; Chen and Flies, 2013; Kaymak et al., 2020; Leone and Powell, 2020). In addition to ICB, CAR-T, and cancer vaccines, many other types of cancer immunotherapies are actively under development. These include oncolytic viruses (Melcher et al., 2021), recombinant cytokines (Li and Lim, 2020; Propper and Balkwill, 2022), gene therapies (Singh et al., 2022), and other forms of immunotherapies such as MAEGI (Wang et al., 2019). Cytotoxic gene therapies have a limitation in that the cancer-killing cargo needs to be delivered into as many cancer cells as possible in order to confer meaningful benefit. On the other hand, immunostimulatory gene therapies can theoretically maintain efficacy even when only a fraction of the tumor cells is successfully transduced, since the resulting immune response would systemically target tumor cells. Several ongoing immune-based cancer gene therapies have been tested, however, these single gene approaches usually suffered from high toxicity or lack of efficacy (Propper and Balkwill, 2022; Hernandez et al., 2022; Mullard, 2021; Melero et al., 2001). We recently demonstrated that CRISPR activation (CRISPRa) systems can be used to overexpress mutant genes thereby tumor neoantigen presentation directly in tumors, driving potent anti-tumor immunity (Wang et al., 2019). A limitation of this approach is the need for potentially complex customized sgRNA libraries targeting patient-specific neoantigen-encoding genes, adding a significant hurdle to clinical translation. An off-the-shelf immunostimulatory gene therapy would therefore be more practical for translation potential.

We reasoned that the challenges faced by current immune-based gene therapies could potentially be overcome by rationally designing an optimal composition of immunostimulatory factors for intratumoral delivery. Outside the cytokines that were well-known to have anti-tumor effects (Propper and Balkwill, 2022), of particular interest were the antigen processing and presentation pathways (Leone et al., 2013; Paschen et al., 2022) and immunomodulatory factors (Demaria et al., 2019), which both play fundamental roles in cancer immunosurveillance and immunotherapy responses, but are not currently targeted by most immunotherapies in clinical use. Nevertheless, to date there has not been a systematic study of the key signals or immunomodulators that can be used to reshape the TME towards a more supportive niche for anti-tumor immunity. We hypothesized that a CRISPR screen focused on antigen presentation and stimulation genes, chemokines, and cytokines could be leveraged to identify, and quantitatively assess, important immunoregulatory molecules for modulating the TME. These targets could then be developed into next-generation immune gene therapies.

As we sought to identify genes that could favorably remodel the TME when ectopically expressed, we adopted a gain-of-function (GOF) CRISPR screen approach. CRISPRa systems, based on the fusion of catalytically dead Cas9 (dCas9) to transcriptional activation elements, enables modular activation and screening of genes at library scale (Konermann et al., 2015). Here, we applied a GOF CRISPRa screen strategy to identify potent immune-activating genes that could then be exogenously delivered as a gene therapy against tumors. Top candidates identified by the screen included *Cd80, Tnfsf14* (encoding LIGHT), *Cxcl10, Tnfsf18* (encoding GITR ligand, GITRL), *Tnfsf9* (encoding 4-1BB ligand, 4-1BBL), and *Ifng* (encoding Interferon gamma). AAV-mediated delivery of these factors induced potent anti-tumor responses, with improved infiltration and activity of cytotoxic T cells. Subsequent optimization of the immunostimulatory gene payload through subtractive screening and additional rationalization further pinpointed a combination of *Ifng, Tnfsf9 (4-1bbl), Il12b* (encoding the IL-12p40 subunit, shared by IL-12 and IL-23) In a humanized setting, we showed that combination therapy with AAV-delivered human transgenes synergized with CAR-T cell therapy against several human solid tumor models.

## Results

### AAV-mediated *in situ* activation of immunoregulatory genes elicits anti-tumor immunity

To test whether multiplexed activation of immunoregulatory genes *in situ* could elicit anti-tumor immunity, we first collected a catalog of 36 immunoregulatory genes involved in antigen processing and presentation (Paschen et al., 2022), innate immune activation (Demaria et al., 2019), as well as cytokines/chemokines (Propper and Balkwill, 2022) and co-stimulatory molecules (Chen and Flies, 2013) for T cell proliferation and migration. This gene set was named the APCM library (for <u>a</u>ntigen <u>p</u>resentation, <u>c</u>ytokine, and co-stimulation <u>m</u>olecules) (**Figure 1A**). AAV is a commonly used delivery system for gene therapies, and two AAV therapies to date have received FDA approval (Kuzmin et al., 2021; Mendell et al., 2017). Key advantages of AAVs include their replication-deficient nature, minimal toxicity, and limited immune responses (Mingozzi and High, 2011). We thus chose AAVs as an *in vivo* delivery system for *in situ* overexpression of immunoregulatory genes. We designed a CRISPRa sgRNA library targeting these immunoregulatory genes and cloned them into an AAV (adeno-associated virus) vector containing a U6 promoter-driven sgRNA cassette and an EF1a promoter-driven MS2-P65-HSF1 (MPH) expression cassette (AAV-CRISPRa vector), generating the AAV-APCM library (**Figure 1B**). To examine the effectiveness of the AAV-APCM library in activating immunoregulatory genes, we first utilized a syngeneic orthotopic triple-negative breast cancer (TNBC) model by transplanting E0771 cells into the mammary fat pad of C57BL/6J mice. We transduced E0771 cells with lentiviral vectors expressing dCas9-VP64 (E0771-dCas9), followed by infection of these cells with AAVs carrying the CRISPRa-APCM sgRNA library (**Figure 1B**). By quantitative RT-PCR, we confirmed the effectiveness of the AAV-APCM library for multiplexed activation of immunoregulatory genes (including *Cd80, Tnfs14, Cxcl10, Tnfsf18, Tnfsf9*, and *Ifng)* in tumor cells (**Figure 1C**). We then tested the therapeutic efficacy of AAV delivering the full library (AAV-APCM) against a TNBC mouse model, established by fat pad injection of E0771-dCas9-VP64. We found pooled multiplexed activation of APCM genes *in situ* showed significant anti-tumor efficacy compared to AAV-Vector or PBS (**Figure 1D**).

**Figure 1.**
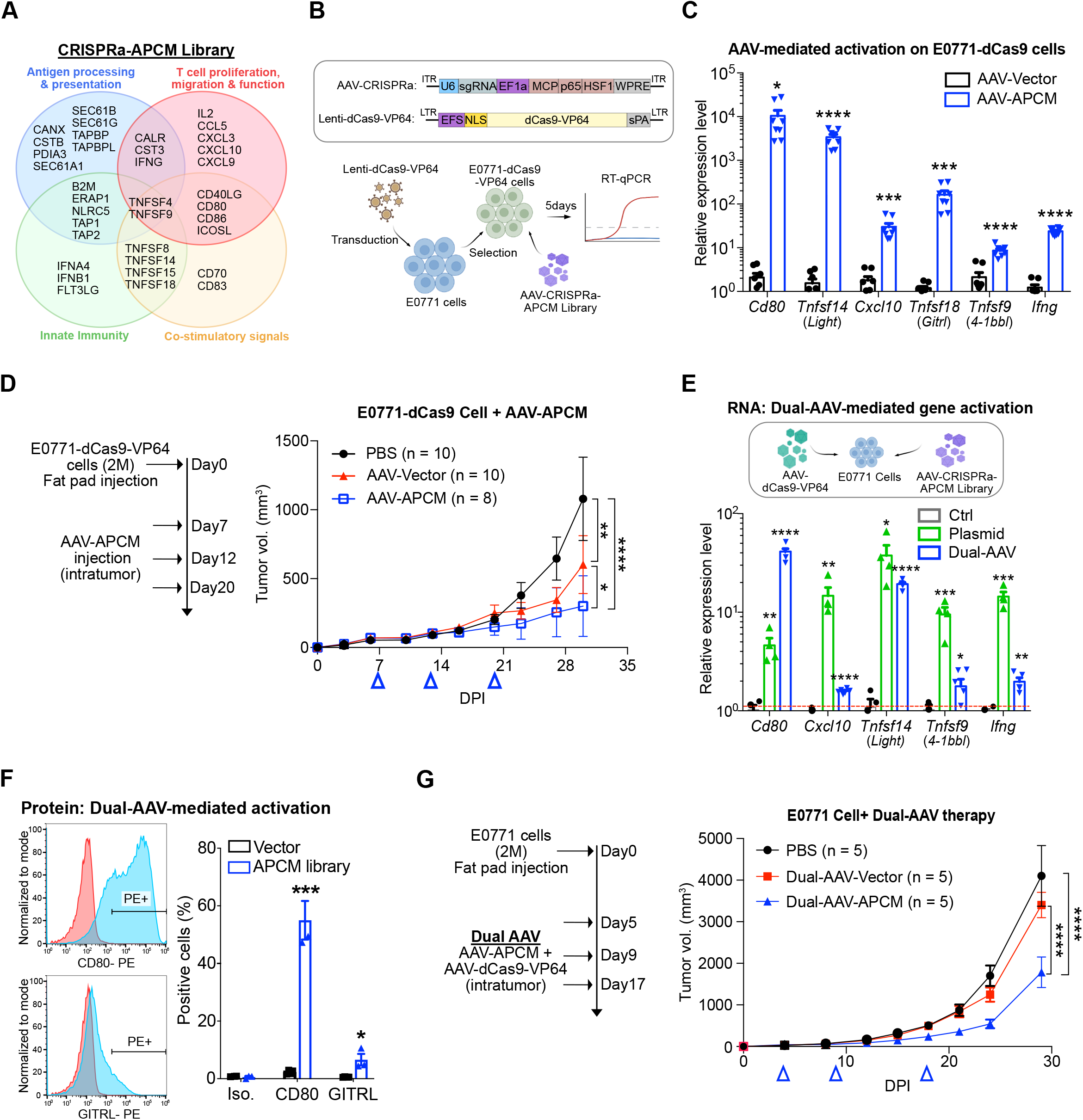
Multiplexed activation of <u>A</u>ntigen <u>P</u>resentation, <u>C</u>ytokine, and Costimulatory <u>M</u>olecules (APCM) as an immunotherapy approach. **(A)**The APCM library targets genes involved in antigen processing and presentation, T cell proliferation and migration that include certain cytokines, as well as stimulatory genes for innate and adaptive immunity. **(B)**Schematic of the experimental approach to characterize gene activation by APCM-gRNAs. dCas9-VP64 overexpressing E0771 cells were infected with AAV-APCM, and the expression of representative targeted genes were analyzed by RT-qPCR at day 5 after infection. qRT-PCR showing AAV-delivered CRISPRa-mediated transcriptional activation of the representative genes including *Cd80, Tnfsf14(Light), Cxcl10, Tnfsf18(Gitrl), Tnfsf9(4-1bbl)*, and *Ifng*. Gene expression was normalized to vector-transduced controls (n = 8 each). **(C)**Tumor growth curves of orthotopic breast tumor formed by E0771-dCas9-VP64 in mice treated by PBS (n = 10 mice), AAV-Vector (n = 10), or AAV-APCM (n = 8) by intratumoral administration at indicated times (blue arrows). **(D)**Dual-AAV infection or plasmid co-transfection mediated CRISPR activation of *Cd80, Cxcl10, Tnfsf14(Light),Tnfsf9(4-1bbl*, and *Ifng*. E0771 cells were infected with the AAV-APCM library or co-transfected with the sgRNA plasmid library and dCas9-VP64 expression plasmids. Expression of representative targeted genes were analyzed by RT-qPCR at day 5 after infection, normalized to vector-transduced controls (n = 8). **(E)**Representative flow cytometry 7 days after dual AAV infection, and quantification of CD80 and TNFSF18 (GITRL) protein on cells infected by dual-AAV-CRISPRa (n = 3). **(F)**Growth curves of E0771 tumors in mice with intratumoral treatment with PBS (n = 5 mice), AAV-dCas9 + AAV-Vector (n = 5), or AAV-dCas9 + AAV-APCM (n = 5) at indicated time points (blue arrows). Data points in this figure are presented as mean ± s.e.m. Statistical significance was asssessed by mutiple unpaired *t*-test (**C**,**E**,**F**) or two-way ANOVA (**D**,**G**). * *p* < 0.05, ** *p* < 0.01, *** *p* < 0.001, **** *p* < 0.0001. Non-significant comparisons not shown.

To evaluate whether the therapeutic effects of AAV-APCM could be observed in tumors that had not been modified to stably express the dCas9-VP64 machinery, we utilized a dual-AAV system composed of a dCas9-VP64-expressing vector (AAV-dCas9-VP64) and the AAV-CRISPRa-APCM-library to deliver the entire CRISPRa system via AAV (**Figure 1E**). By *in vitro* co-infection of E0771 cells with the dual-AAV system, we first confirmed that targeted genes such as *Cd80, Cxcl10, Tnfsf14, Tnfsf9* and *Ifng* could be successfully activated using qRT-PCR (**Figure 1E**). We also confirmed gene activation on the protein level by flow cytometry (**Figure 1F**). To test the therapeutic efficacy of the dual-AAV delivered APCM CRISPRa library, we co-injected both AAV-dCas9 and AAV-APCM into established E0771 breast tumors. Similarly, we observed significant anti-tumor effects with the dual-AAV delivery (**Figure 1G**), and in this case, the AAV-vector treatment had a negligible effect on tumor growth (**Figure 1G**), confirming the therapeutic activity of forced activation of APCM genes by CRISPRa.

### A CRISPR activation screen identifies immune-activating molecules in metastatic foci

While AAV-APCM demonstrated anti-tumor activity, we sought to optimize and distill the original APCM library into a smaller set of potent components. We thus performed an *in vivo* CRISPR activation screen in the setting of an immunocompetent host. We first established dCas9-VP64 and MPH expressing E0771 cells by lentiviral infection (E0771-dCas9-MPH) and then transduced them with a lentiviral version of the APCM CRISPRa sgRNA library (Methods) (**Figure 2A**). After confirming the successful activation of several representative genes in the APCM library-transduced cells by flow cytometry (**Figure 2B**), we intravenously injected the library-transduced cells into the immunocompetent C57BL/6J mice, leading to the eventual formation of numerous independent metastatic lung foci. We then harvested all endpoint lung lobes with metastases to analyze the enriched or depleted sgRNAs in immune-edited metastatic tumors *vs*. the *in vitro* pre-injection cell pool (**Figure 2A&C**). To compare the sgRNA abundances in lung metastatic tumors with the *in vitro* pre-injection cell pool, we first aggregated the data to the gene level, then performed linear regression using the data from the entire APCM library. Several genes were outside the 95% confidence interval of the linear regression (**Figure 2C**). The genes below the linear regression line (indicating relative depletion in tumors) included *Cd80, Tnfsf14/Light, Cxcl10, Tnfsf18/Gitrl, Tnfsf9/4-1bbl*, and *Ifng*. In sum, the CRISPRa screen identified several candidate factors that led to reduced lung metastatic potential when overexpressed in TNBC cells, suggesting that immune responses were mounted in response to these factors (**Figure 2C**).

**Figure 2.**
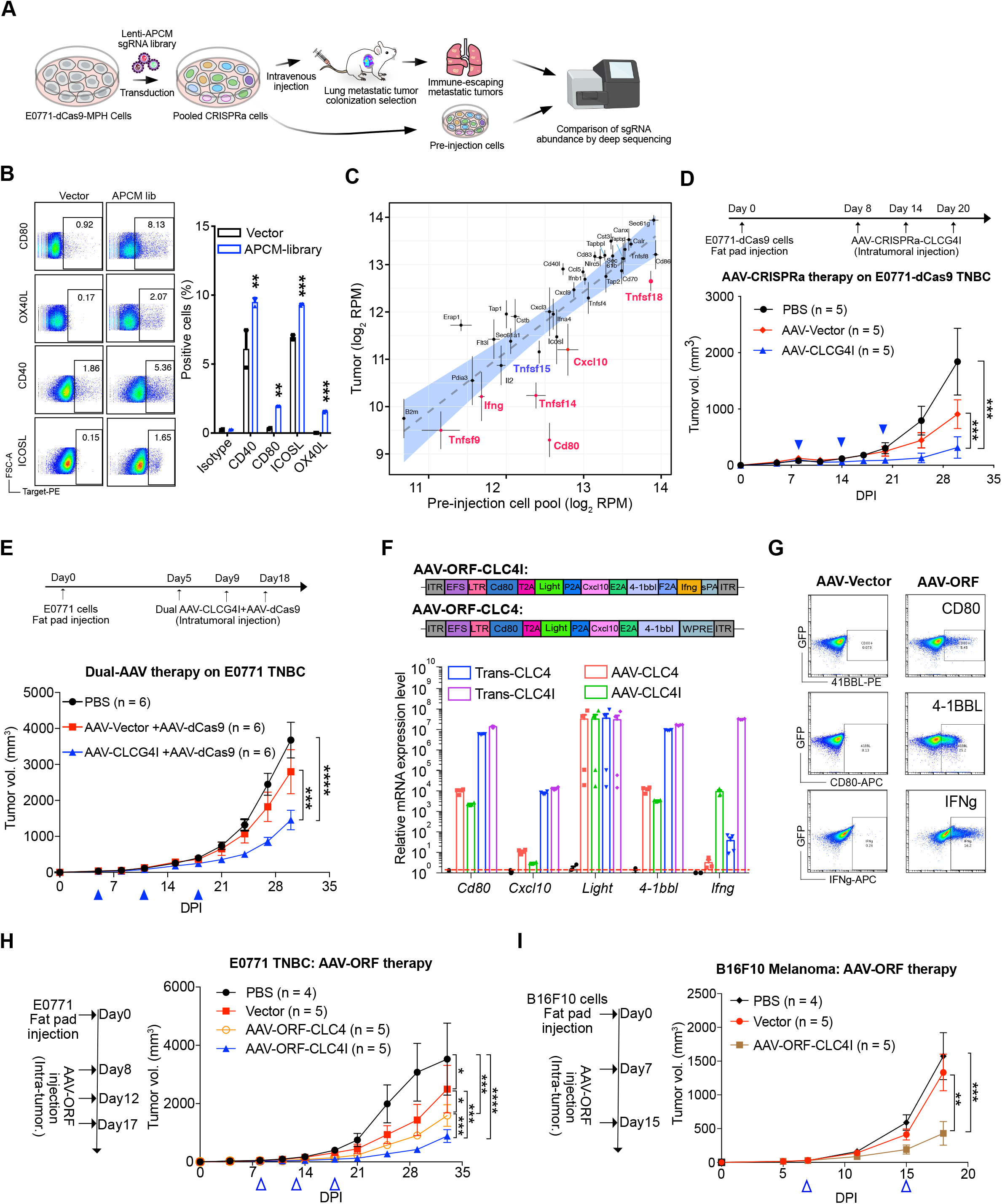
An *in vivo* CRISPR activation screen reveals key TME immunomodulators. **(A)**Schematic of the APCM CRISPRa screen design. The E0771 breast cancer cells were transduced with lentiviruses carrying the APCM sgRNA library, then intravenously injected into immunocompetent C57BL/6J mice. The lung-metastatic tumors were collected for sgRNA enrichment and depletion analysis in comparison to the pre-injected cells. **(B)**Flow cytometry of representative co-stimulatory molecules such as CD80, CD40, OX40L, and ICOSL in APCM library-transduced E0771 cells. **(C)**Average APCM gene representation across metastatic tumors in C57BL/6J mice (n = 14) was plotted against average APCM gene representation in the cultured cell pool (n = 2). The linear regression of gene-level abundance is shown, with 95% confidence intervals. Select genes lying underneath the regression line (suggestive of *in vivo* depletion from immunoediting pressure) are colored. **(D)**Growth curves of E0771 tumors after intratumoral treatment with PBS (n = 5), AAV-Vector (n = 5), AAV-CLCG4I (n = 5). AAV-CLCG4I: AAV-CRISPRa vector expressing *Cd80, Cxcl10, Light, Gitrl, 4-1bbl*, and *Ifng* (CLCG4I). **(E)**Growth curves of E0771 tumors after intratumoral treatment with PBS (n = 6), Dual-AAV-Vector (n = 6), Dual-AAV-CLCG4I (n = 6). **(F)**RT-qPCR analysis of AAV-mediated ectopic gene delivery with 2A-spaced polycistronic open reading frames (AAV-ORF). AAV-ORF-CLC4I: *Cd80+Light+Cxcl10+4-1bbl*+*Ifng*. AAV-ORF-CLC4: *Cd80+ Light+ Cxcl10+4-1bbl*. **(G)**Representative flow plots detailing the ectopic expression of CD80, 4-1BBL, and IFNg in E0771 cells at the protein level with AAV-ORF treatment *in vitro*. **(H)**Growth curves of E0771 tumors treated with PBS (n = 4), AAV-Vector (n = 5), AAV-ORF expression of *Cd80, Cxcl10, Light*, and *4-1bbl* (AAV-ORF-CLC4, n = 5), or AAV-ORF of *Cd80, Cxcl10, Light, 4-1bbl*, and *Ifng* (AAV-ORF-CLC4I, n = 5). **(I)**Therapeutic effects of intratumorally delivered PBS (n = 4), AAV-Vector (n = 5), or AAV-ORF-CLC4I (n = 5) on B16F10 melanoma tumors. AAV-ORF-CLC4I:*Cd80+Cxcl10+Light+4-1bbl*+*Ifng*. Data points in this figure are presented as mean ± s.e.m. Statistical significance was assessed by multiple unpaired *t*-test (**B**) or two-way ANOVA (**D-G**). * *p* < 0.05, ** *p* < 0.01, *** *p* < 0.001, **** *p* < 0.0001. Non-significant comparisons not shown.

We then investigated whether *in situ* overexpression of the identified genes by AAV could induce anti-tumor immune responses against established tumors. To test the therapeutic effects of the hyper-expression of these genes, we pooled the CRISPRa sgRNAs targeting *Cd80, Cxcl10, Light, Gitrl, 4-1bbl* and *Ifng* (hereafter CLCG4I) and made an AAV minipool for multiplexed activation of these genes. We examined the therapeutic effects of AAV-delivered CLCG4I on syngeneic breast tumors formed by E0771-dCas9 cells in C57BL/6J mice, finding that activation of *Cd80, Light, Cxcl10, Gitrl, 4-1bbl*, and *Ifng* in the TME showed strong anti-tumor effects compared to either PBS or AAV-Vector (**Figure 2D**). Moreover, the therapeutic effect was recapitulated using the dual-AAV system for multiplexed activation of the above immune regulatory genes in unmodified E0771 tumors (**Figure 2E**).

Aside from CRISPRa activation, gene open-reading-frame (ORF) is also a reliable way to achieve gene overexpression, which also doesn’t involve CRISPR machinery. Another advantage of the ORF delivery is that multiple ORFs can be expressed in a single AAV vector given that the transgenes are within the AAV packaging limit, thereby further simplifying the manufacturing process. Having the immune-activating genes identified by the CRISPRa screen, we then sought to perform a parallel validation of this multiplex gene activation immunotherapy using a simplified ORF viral vector system. To do so, we designed an AAV with an EFS promoter driving the expression of a 2A-spaced polycistronic ORF (all-in-one ORF), for multiplexed overexpression of the immune-activating molecules identified in the screen (**Figure 2F**). We designed two AAV-all-in-one ORFs with or without *Ifng* (AAV-CLC4: *Cd80* + *Light* + *Cxcl10* + *4-1bbl*, and AAV-CLC4I: *Cd80* + *Light* + *Cxcl10* + *4-1bbl* + *Infg*) (**Figure 2F**). Through both *in vitro* plasmid transfection and AAV infection assay on E0771 cells, we confirmed the ORF-mediated multiplexed expression of *Cd80, Light, Cxcl10, 4-1bbl*, and *Ifng* at the RNA level by qRT-PCR (**Figure 2F**). We also confirmed AAV-mediated expression of CD80, 4-1BBL and IFNg at the protein level by flow cytometry (**Figure 2G**).

Next, we evaluated the anti-tumor effects of multiplexed *in situ* overexpression of AAV-CLC4 or AAV-CLC4I in a syngeneic orthotopic TNBC model (E0771 in C57BL/6J mice). We found that both AAV-CLC4 and AAV-CLC4I showed robust anti-tumor effects compared to PBS or AAV-Vector (**Figure 2H)**. These data indicate that multiplexed immune gene activation can have significant anti-tumor effects, with both AAV-ORF overexpression and AAV-CRISPRa approaches demonstrating significant therapeutic efficacy. We also noted that AAV-CLC4I had significantly stronger anti-tumor effects than AAV-CLC4 (**Figure 2H)**, indicating the potency of exogenously delivered IFN-γ in driving anti-tumor responses. To test the broader utility of AAV-CLC4I as a therapeutic approach, we further tested it in a syngeneic melanoma model (B16F10 in C57BL/6J), and again found that AAV-mediated multiplexed expression of *Cd80, Light, Cxcl10, 4-1bbl*, and *Ifng* induced robust anti-tumor effects (**Figure 2I**). Thus, from both AAV-CRISPRa minipool or all-in-one AAV-ORF strategies, multiplex activation of APCM genes showed robust anti-tumor effect in different tumor models.

### Optimization of the immune-activating gene combinations for robust anti-tumor responses

To examine whether the expression of all six identified genes were required for driving anti-tumor immunity, we further performed “minus-one” subtractive efficacy screening in the E0771 TNBC model, by removing one gene at a time from the six-gene combination (AAV-CRISPRa CLCG4I-minus-One). In all cases, we observed significant anti-tumor effects of the multiplexed activation therapies in comparison to PBS or AAV-Vector (**Figure 3A**), regardless of which individual immunoregulatory factor was excluded. By continuing observation of the tumor growth over a longer period of time, we started to observe the efficacy differences (**Figure 3B**). When compared to the complete CLCG4I AAV-CRISPRa minipool combination, the removal of *Ifng* or *41bbl* among the groups most dramatically decreased the anti-tumor effects (**Figure 3B**). In contrast, the removal of *Light, Gitrl, Cxcl10* or *Cd80* from the pool of CLCG4I only modestly affect the anti-tumor activity when compared to CLCG4I (these groups have numerically slightly faster tumor growth curves than CLCG4I but not significant) (**Figure 3B**). Thus, the CRISPR activation screen and the “minus-one” efficacy testing highlighted *Ifng* and *4-1bbl* as potent factors for eliciting strong anti-tumor responses.

**Figure 3.**
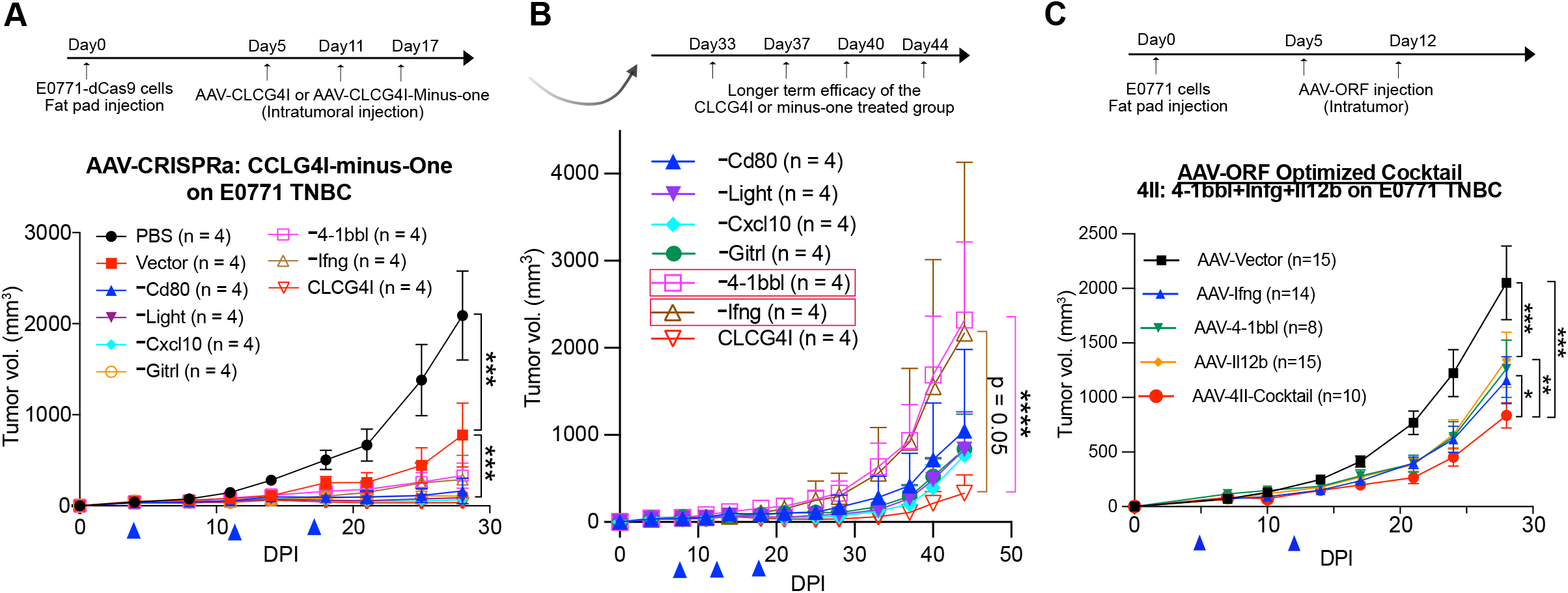
Rationalization of immunostimulatory gene combinations for robust anti-tumor responses. **(A)**Optimization of the CLCG4I gene subtractive “minus one” screening. Growth curves of E0771 tumors treated by PBS (n = 4), AAV-Vector (n = 4), AAV-CLCG4I (n = 4), or AAV-CLCG4I-minus-One (n = 4 for each condition). AAV-CLCG4I: CRISPRa guide RNA minipool targeting *Cd80, Cxcl10, Light, Gitrl, 4-1bbl*, and *Ifng*. **(B)**Long-term growth curves of E0771 tumors treated by AAV-CLCG4I (n = 4), or AAV-CLCG4I-minus-One (n = 4 for each condition), extended time points of the same experiment as (**A**). **(C)**Growth curves of E0771 tumors treated with AAV-Vector (n = 15), AAV-Ifng (n = 14), AAV-4-1bbl (n = 8), AAV-Il12b (n = 15), or a cocktail of AAV-ORFs containing Ifng+41bbl+Il12b (n = 10). IL12b encodes Il12p40 subunits shared by Il12 and Il23. Data points in this figure are presented as mean ± s.e.m. Statistical significance was assessed by two-way ANOVA. * *p* < 0.05, ** *p* < 0.01, *** *p* < 0.001, **** *p* < 0.0001. Non-significant comparisons not shown.

According to a recent study, T cells engineered with the IL-12β p40 (*Il12b*) showed improved proliferation and antitumor capacity, because the antigen-recognizing T cells could only upregulate IL-23α, but still need IL-12β to form the key immunoproliferative cytokine IL23 (Ma et al., 2020). We therefore tested the anti-tumor effect of AAV delivered IL-12β (AAV-Il12b), and found that AAV-Il12b treatment could achieve a similar anti-tumor response as our top two candidate genes, *Ifng* and *4-1bbl* (**Figure 3C)**. We then tested the anti-tumor effects of combined expression of *<u>4-</u>1bbl, <u>I</u>fng*, and *<u>I</u>l12b* (AAV-4II) in a syngeneic E0771 TNBC model. According to the tumor growth curves, the combinatorial expression of *4-1bbl, Ifng*, and *Il12b* (AAV-4II) showed significantly more potent anti-tumor effects compared to AAV-4-1bbl, AAV-Ifng or AAV-Il12b alone (**Figure 3C**). Nevertheless, the intratumoral treatments with single-gene AAVs did show significant anti-tumor effects compared to AAV-Vector control (**Figure 3C**). Taken together, these data demonstrate that the AAV-mediated delivery of *4-1bbl, Ifng*, and *Il12b*, either alone or in combination, can elicit significant anti-tumor immune response.

### AAV-mediated expression of immunoregulatory genes reshaped the TME

Since the combinatorial overexpression of *4-1bbl, Ifng*, and *Il12b* by AAV elicited significant anti-tumor responses, we analyzed how their *in situ* expression affects the immune context and status of TME by flow cytometry (**Figure 4**). We found only the combination of *Ifng*+*Il12b*+*4-1bbl* (AAV-4II), but neither *Ifng* nor *Il12b* alone, significantly increased the infiltration of CD45^+^ immune cells in the tumor compared to AAV-Vector (**Figure 4A**). Furthermore, the combination of *Ifng*+*Il12b*+*4-1bbl* also induced significantly higher levels of CD45^+^ immune cell compared to the treatment of Ifng or Il12b alone (**Figure 4A**). Further characterizing the changes of sub-populations among tumor-infiltrating immune cells, we found that the combination of *Ifng*+*Il12b*+*4-1bbl*, but not *Ifng* or *Il12b* alone, elicited significantly higher levels of tumor-infiltrating CD8^+^ T cells (**Figure 4B**), including IFN-γ-producing CD8^+^ T cells (**Figure 4C**) but not PD1^+^CD8^+^ T cells (**Figure 4D)**. Additionally, we observed significantly higher percentages of tissue-resident memory CD8^+^ T cells (defined by CD45^+^CD3^+^CD8^+^CD69^+^CD103^+^) after *Ifng*+*Il12b*+*4-1bbl* treatment compared either to AAV-Vector or single *Il12b* treatment, though this effect was also observed with *Ifng* treatment alone (**Figure 4E**). In these groups, we did not observe any significant changes in total CD4^+^ T cells (**Figure 4F**), IFN-γ producing CD4^+^ T cells (**Figure 4G**), or tissue-resident memory CD4^+^ T cells (**Figure 4H**). As for the innate immune cells, there is no obvious difference between these groups in neutrophils (**Figure 4I**), monocytes (**Figure 4J**), dendritic cells (**Figure 4K**) or macrophages (**Figure 4L**). These results suggested that the multiplexed expression of *Ifng*+*Il12b*+*4-1bbl* enhanced infiltration of total CD8^+^ T cells, including IFN-γ producing CD8^+^ T cells and tissue resident CD8^+^ T cells in the TME.

**Figure 4.**
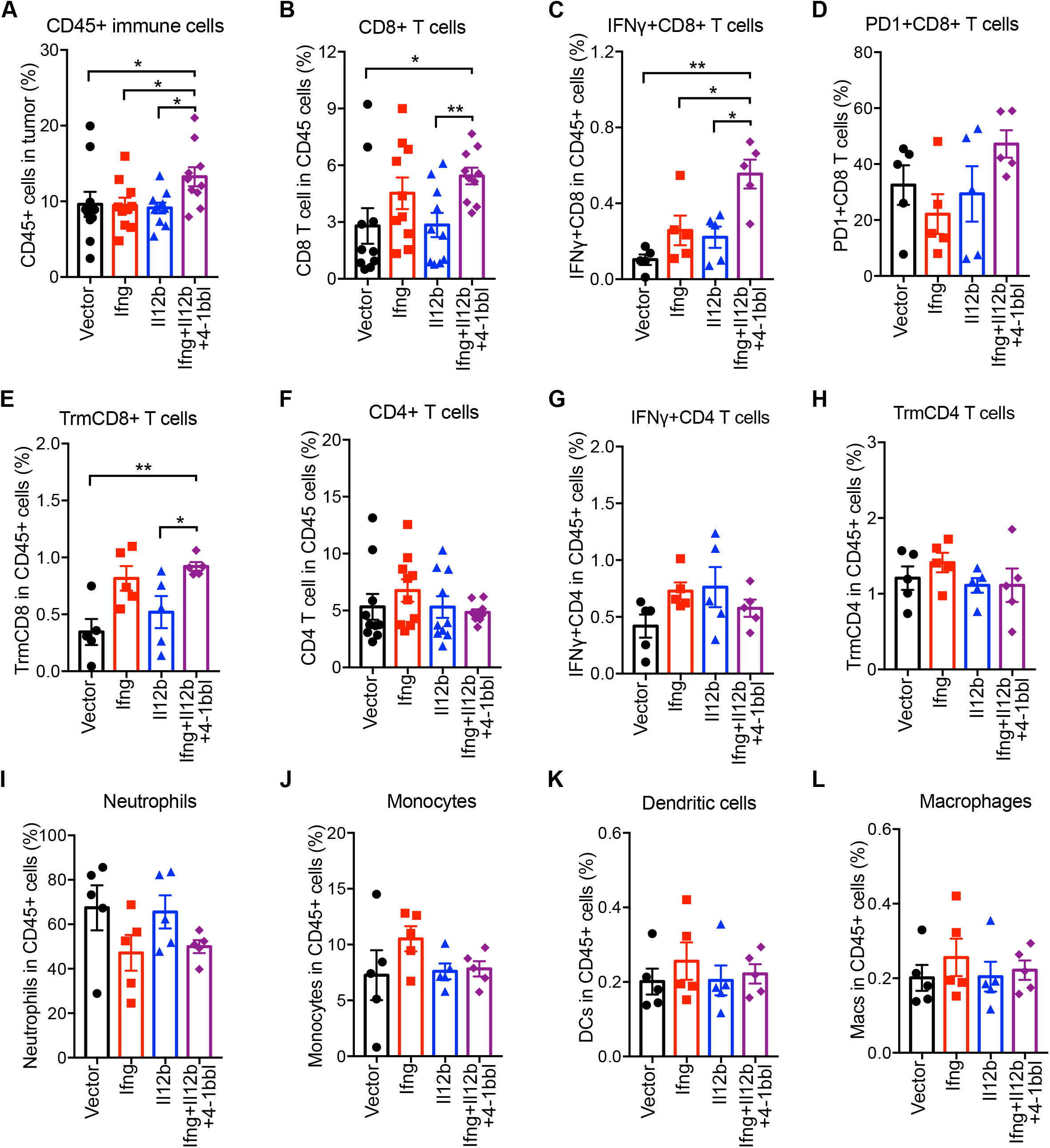
AAV-ORF delivery of immunostimulatory molecules promotes immune infiltration into the TME. (**A**) Flow cytometry quantification of CD45^+^ immune cells in the E0771 tumor microenvironment at DPI = 28. Percentage of CD45^+^ immune cells out of total cells in tumors treated with AAV-Vector, AAV-Ifng, AAV-Il12b, or AAV-Ifng+Il12b+4-1bbl. (**B-E**) Percentage of total CD8^+^ T cells (**B**), IFN-γ^+^CD8^+^ T cells (**C**), PD1^+^CD8^+^ T cells (**D**), and tissue-resident memory CD8^+^ T cells (**E**), among tumor-infiltrating CD45^+^ immune cells. (**F-H**) Percentage of total CD4^+^ T cells (**F**), IFN-γ^+^CD4^+^ T cells (**G**), and tissue-resident memory CD4^+^ T cells (**H**) among tumor-infiltrating CD45^+^ immune cells. (**I-L**) Percentage of neutrophils (**I**), monocytes (**J**), dendritic cells (**K**), and macrophages (**L**) among tumor-infiltrating CD45^+^ immune cells. Data points in this figure are presented as mean ± s.e.m. Statistical significance was assessed by Mann– Whitney test. * *p* < 0.05, ** *p* < 0.01. Non-significant comparisons not shown.

### AAV-mediated immune gene therapy synergizes with CAR-T therapy against solid tumors

We wondered whether the TME remodeling using AAV-delivered immunoregulatory genes have mutual enhancement together with T cell therapies against solid tumors. We first generated HER2-targeting CAR-T cells by transducing freshly isolated, anti-CD3/anti-CD28 activated peripheral blood mononuclear cells (PBMCs) with lentiviral EFS-scFv(Herceptin)-CD8TM-41BBL-CD3zeta-T2A-Puro-WPRE (**Figure S1A&B**). As a proof of principle, we first utilized an *in vitro* co-culture assay to investigate whether overexpression of the top immunoregulators could improve tumor cell killing by CAR-T cells. After transducing HER2-expressing MCF7 cells with lentiviruses for the expression of luciferase as well as human *IFNG, 4-1BBL, IL12B*, or *CD80*, we co-cultured these tumor cells with HER2-CAR-T cells. The *in vitro* overexpression of IFNg, 4-1BBL, IL12B, or CD80 led to significantly enhanced killing of cancer cells after 16 hours of coculture across two effector : target (E:T) ratios (**Figure 5A**).

**Figure 5.**
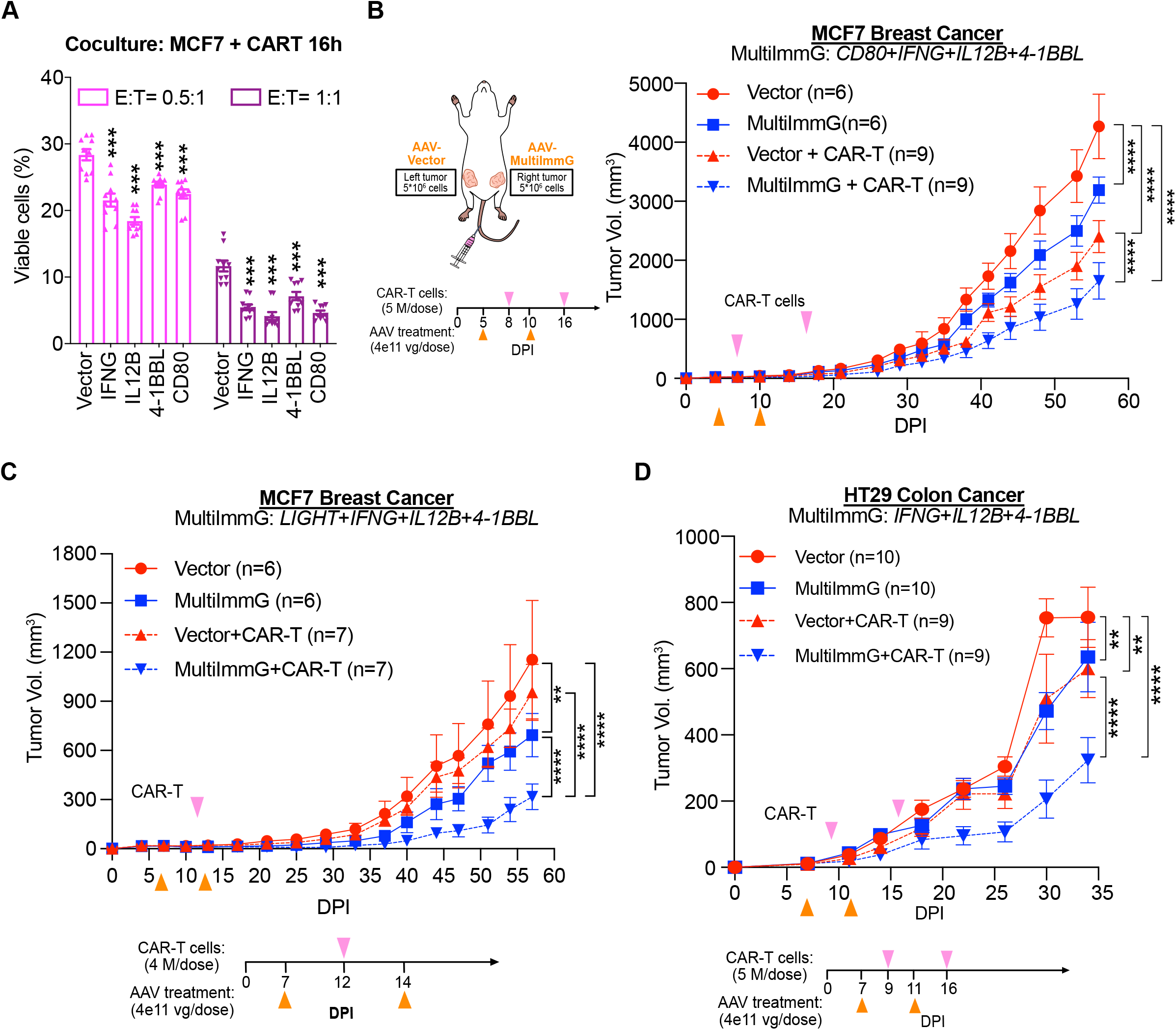
AAV delivery of immunostimulatory molecules synergizes with CAR-T cell therapy in human cancer models. **(A)**MCF7 human breast cancer cells transduced with human *CD80, IFNG, IL12B*, or *4-1BBL* expressing lentiviruses were co-cultured with HER2 CAR-T cells. After 16 hours of coculture, the remaining viability of MCF7 cells were measured (n = 10). Multiple unpaired *t*-test, *** *p* < 0.001. **(B)***In vivo* efficacy of HER2 CAR-T cells and an AAV-MultiImmG combination (CD80+IFNG+IL12B+4-1BBL) in a dual-tumor MCF7 human breast cancer model. Mice were intratumorally injected with AAV-Vector at the left site and AAV-MultiImmG at the right site on the indicated days. CAR-T cells were intravenously injected on the indicated days. **(C)**The combination therapeutic effects with an AAV-MultiImmG combination (LIGHT+INFG+IL12B+4-1BBL) therapy and CAR-T in the human breast tumors formed by injecting MCF7 into NSG mice. Mice were intratumorally injected with AAV-Vector at the left site or AAV-MultiImmG at the right site on the indicated days. CAR-T cells were subsequently intravenously injected on indicated days. **(D)***In vivo* efficacy of HER2 CAR-T cells and an AAV-MultiImmG combination (IFNG+IL12B+4-1BBL) in a dual-tumor HT29 human colon cancer model. Mice were intratumorally injected with AAV-Vector at the left site and AAV-MultiImmG at the right site on the indicated days. CAR-T cells were intravenously injected on the indicated days. Data points in this figure are presented as mean ± s.e.m. Statistical significance was assessed by two-way ANOVA (**B-D**). * *p* < 0.05, ** *p* < 0.01, *** *p* < 0.001, **** *p* < 0.0001. Non-significant comparisons not shown. **See also: Figure S1**

To test the potential synergy of therapeutic efficacy between AAV-mediated endogenous gene activation (AAV-MultiImmG) and CAR-T in a human gene setting *in vivo*, we utilized a dual tumor model by injecting the human cancer cells into both sides of the mammary fat pads or flanks of NSG mice (**Figure 5B**), with an advantage that both tumors shared the same source of CAR-T cells to allow better control and comparison of efficacy. We then intratumorally injected AAV-Vector, the AAV-MultiImmG combination into the left and right site tumor respectively, followed by CAR-T infusion. In a xenograft model of MCF7 breast cancer, the HER2 CAR-T treatment itself has significant efficacy (**Figure 5B**); the treatment with AAV-MultiImmG (*CD80+IFNG+4-1BBL+IL12B*) also has significant efficacy (**Figure 5B**); while the combination therapy with AAV-MultiImmG and HER2 CAR-T cells have strong synergy in eliciting significantly better anti-tumor effects than either alone (**Figure 5B**). In another independent experiment, we tested a different combination of APCM genes (*LIGHT+IFNG+4-1BBL+IL12B*) against the same MCF7 breast cancer (**Figure 5C**). AAV-MultiImmG with this combination again showed significant anti-tumor efficacy alone, and importantly, significant synergy with HER2 CAR-T cells (**Figure 5C**).

Furthermore, we evaluated the three-gene combination of human *IFNG+4-1BBL+IL12B*. In a MCF7 xenograft breast model, we found the tumors in all groups were inhibited to a baseline level when treated with high titer AAVs (**Figure S1C**). We subsequently re-challenged these mice with tumor cells again and then treated the mice with AAV-MultiImmG/AAV-Vector and CAR-T cells (**Figure S1C**). We found the combination therapy with CAR-T cells and AAV-MultiImmG expressing *IFNG+4-1BBL+IL12B* showed stronger anti-tumor efficacy against the relapse of breast tumors than AAV-vector, AAV-MultiImmG alone, or CAR-T cells alone (**Figure S1C**). We finally tested the AAV-MultiImmG (*IFNG+4-1BBL+IL12B* combo) plus CAR-T cell therapy combination in an independent HT29 colon cancer model, again by dual tumor inoculation, subcutaneously in this setting. We again observed that this AAV APCM gene combination showed significant anti-tumor efficacy alone, and importantly, significant synergy with HER2 CAR-T cells against HT29 colon cancer *in vivo* (**Figure 5D**). These data demonstrate that multiplexed *in situ* expression of several combinations (*CD80+IFNG+4-1BBL+IL12B* combo, *LIGHT+IFNG+4-1BBL+IL12B* combo and *IFNG+4-1BBL+IL12B* combo) all have significant synergy with CAR-T cells against solid tumors in human breast and/or colon cancer models.

## Discussion

The recognition and killing of cancer cells by T lymphocytes is essential for the clinical success of cancer immunotherapies (Schumacher and Schreiber, 2015)3/14/23 6:17:00 PM. However, the immunosuppressive TME formed by solid tumors often leads to T cell exclusion and dysfunction through multiple mechanisms (Joyce and Fearon, 2015). We previously showed that AAV-based CRISPRa can serve as a new form of TME-remodeling gene therapy, where *in situ* delivery of AAV-CRISPRa resulted in magnification of the “non-self” signals of the tumor and elicited potent anti-tumor immunity (Wang et al., 2019). Besides the direct targeting of mutant genes, the expression of immune regulatory molecules for antigen presentation, innate immunity, cytokines, as well as the co-stimulatory receptors also play a pivotal role in the status and fate of T cells (Chen and Flies, 2013). In this study, we employ an *in vivo* GOF screening strategy in a metastatic cancer context with immune pressure, to identify the key immunomodulators whose *in situ* expression could reshape TME toward more supportive niche for anti-tumor immunity. With this approach, we identified several key modulators including *Cd80, Tnfsf14, Cxcl10, Tnfsf18, Tnfsf9, Ifng* that can remodel the TME when delivered through AAVs. These are highly conserved common genes and can be targeted either alone or in combinations as a set of universal immune gene therapy.

In parallel, we tested both an AAV-delivered CRISPRa system and an AAV-delivered ORF expression system for the *in situ* expression of the identified immunomodulators, demonstrating that both systems could elicit strong antitumor effects. The advantage of AAV-CRISPRa-APCM is that it can rapidly customize compositions or cocktails of immune gene therapies with ease, simply by generating a pool of effective sgRNAs to be delivered together with the shared CRISPRa machinery. The advantage of AAV-ORF-APCM on the other hand, is that it can encode multiple transgenes into one AAV vector for delivery (as in the scenario here). Both AAV-CRISPRa-APCM and AAV-ORF APCM drastically simplified manufacturing as compared to MAEGI, with AAV-ORF having the most simplicity for manufacturing. As compared to traditional single-gene cancer gene therapy, the APCM screened and rationalized combos have the advantage of being less biased and not dependent on a single gene. APCM and the immunotherapeutic combinations here scale between p-MAEGI and traditional single-gene therapy, balancing the anti-tumor efficacy and manufacturability. Finally, *ex vivo*-expanded adoptive cell therapies with naturally occurring or genetically engineered T cells have achieved limited success against solid tumors, largely due to the suppressive TME (Morotti et al., 2021). We showed that the combinatorial use of CAR-T and TME-remodeling immunoregulators delivered by AAV can substantially enhance each other’s anti-tumor efficacy, leading to significant synergistic effects. A limitation to this approach is the need for intratumoral AAV delivery.

In summary, we leveraged a CRISPRa screen as an approach to identify genes as cargos for cancer immune gene therapy. We selected and validated several key immuno-modulators and their combinations that can be used to reshape the TME towards a more supportive niche for anti-tumor immunity. The resultant immunostimulatory genes enabled the rapid and rational design of gene therapy combinations that could reshape the suppressive TME, with potent anti-tumor efficacy. In human cancer models with adoptive CAR-T therapy, the data in solid tumor models here showed that the combination activation therapies delivered by AAV can synergize with CAR-T cell therapy, achieving significantly boosted therapeutic efficacy compared to either CAR-T cell or AAV gene therapy alone.

## Author Contributions

Conceptualization: GW, SC. Design: GW, RC, SC. Experiment lead: GW, FZ. Analytic lead: RC. Experiment assistance and support: EH, LZ, QH. Manuscript prep: GW, FZ, RC, SC. Supervision and funding: SC.

## Acknowledgments

We thank all members in Chen laboratory, as well as various colleagues in Yale Genetics, SBI, CSBC, MCGD, Immunobiology, BBS, YCC, YSCC, and CBDS for assistance and/or discussions. We thank various Yale Core Facilities such as YCGA, HPC, WCAC, KBRL for technical support.

S.C. is supported by NIH/NCI/NIDA (DP2CA238295, R01CA231112, R33CA225498, RF1DA048811), DoD (W81XWH-17-1-0235, W81XWH-20-1-0072, W81XWH-21-1-0514), Damon Runyon Dale Frey Award (DFS-13-15), Melanoma Research Alliance (412806, 16-003524), Cancer Research Institute (CLIP), AACR (17-20-01-CHEN), The V Foundation (V2017-022), Alliance for Cancer Gene Therapy, Sontag Foundation (DSA), Pershing Square Sohn Cancer Research Alliance, Dexter Lu, Ludwig Family Foundation, Blavatnik Family Foundation, and Chenevert Family Foundation. GW is supported by CRI Irvington and RJ Anderson Postdoctoral Fellowships. RC is supported by NIH MSTP training grant (T32GM007205) and NRSA fellowship (F30CA250249).

## Data and material availability

All data generated or analyzed during this study are included in this article and its supplementary information files. Source data and statistics are provided in an excel file of **Source data and statistics**. Processed data for genomic sequencing and gene expression are provided as processed quantifications in **Supplementary Datasets**. Genomic sequencing raw data are being deposited to NIH Sequence Read Archive (SRA) and/or Gene Expression Omnibus (GEO), with pending accession numbers. Data, codes and materials that support the findings of this research are available from the corresponding author upon reasonable request to the academic community.

## Methods

### Institutional Approval

This study has received institutional regulatory approval. All biosafety work was performed under the guidelines of Yale Environment, Health and Safety (EHS) Committee with an approved protocol (Chen-rDNA-15-45; Chen-rDNA-18-45). All animal work was performed under the guidelines of Yale University Institutional Animal Care and Use Committee (IACUC) with approved protocols (Chen 2015-20068; 2018-20068; 2021-20068). All human sample work was performed under the guidelines of Yale University Institutional Review Board (IRB) with an approved protocol (HIC#2000020784).

### Design and cloning of CRISPR activation library

The well-recognized antigen-presenting genes, immune costimulatory molecules, chemokines and cytokines were chosen, and named as APCM library. The sgRNA library targeting the above genes were designed using sgRNA Designer of Broad Institute (https://broad.io/crispick, targeting the window within 200 bp upstream of each TSS and filtered for GC content > 25%). The oligos of sgRNAs were synthesized in IDT technology, annealed, and cloned into lentiviral vector of CRISPR activation (U6-sgRNA(ms2)-EFS-Puro-WPRE). The lentiviral sgRNA library was expanded by electroporation as previous described. An estimated library coverage of >100× (>100 colonies per sgRNA) was achieved in electroporation, and the coverage of sgRNAs was subsequently sequenced and verified by Illumina sequencing.

### Lentivirus production

For lentivirus production, 20 μg of cis-lentiviral vector like lenti-EF1a-NLS-dCas9-VP64-P2A-Blast-WPRE, lenti-EF1a-MS2-p65-HSF1-P2A-Hygro-WPRE, target sgRNA or library in lenti-U6-sgRNA(ms2)-EFS-Puro-WPRE, together with 10 μg of pMD2.G and 15 μg of psPAX2 were co-transfected into HEK293FT cells in 150 mm-dish at 80-90% confluency using 130 μg polyethyleneimine (PEI) as the transfection reagent. 6-12 hours later, the media was replaced by fresh DMEM+10%FBS. Virus supernatant was collected 48 h and 72 h post-transfection, centrifuged at 1,500 g for 10 min and passed through 0.45-μm filter to remove the cell debris; aliquoted and stored at -80°C. Library virus was titrated by infecting E0771 cells followed by the selection under 5 μg/ml puromycin.

### Generation of mAPCM library- and vector-transduced cells

E0771 cells stably expressing dCas9-VP64 (E0771-dCas9) were generated by transducing E0771 cells with lentiviral EF1a-NLS-dCas9-VP64-2A-Blast-WPRE, followed by 4-7 days of selection under 10-20 μg/ml blasticidin. E0771-dCas9 cells were further transfected with lentiviral EF1a-MS2–p65–HSF1-2A-Hygro-WPRE and selected under 500 μg/ml hygromycin to generate dCas9-VP64-MPH expressing E0771 cells (E0771-dCas9-MPH). To generate library cell pool was generated by infecting the E0771-dCas9-MPH cells with CRISPRa sgRNA library-containing lentiviruses. To guarantee enough coverage, 2 × 10^7^ cells of E0771-dCas9-MPH cells were infected with lentiviral CRISPRa sgRNA library of mAPCM at a calculated MOI of 0.3 with a minimal representation of > 200× transduced cells per sgRNA as previously described. Library or vector infected cells were cultured at 37°C more than 24 h before selecting under 3-5 μg/ml puromycin containing media for 2-3 days, and the transduced cells were cultured for more than 5 days before use.

### Mice

Female mice at the ages of 5-12 weeks of age were used for the study unless otherwise specified. All animals were housed in standard individually ventilated, pathogen-free conditions, with light cycle of 12h:12h, room temperature of 21-23°C, and relative humidity of 40-60%. Sample size (i.e. number of mice per treatment group) was determined on similar tumor models in the field. When a cohort of animals receive multiple treatments, animals were randomized by 1) randomly assigning littermates to different groups prior to treatment, maximizing the evenness or representation of mice from different cages in each group, 2) group animals prior to treatment so that each group has even distribution of initial tumor sizes; and/or 3) random assignment of mice to minimize the effect of litter, and small differences in age, cage, or housing position.

### Genomic DNA extraction from cells and mouse tissues

For gDNA extraction, 50-200 mg of frozen ground tissue were resuspended in 6 ml of Lysis Buffer (50 mM Tris, 50 mM EDTA, 1% SDS, pH 8.0) in a 15 ml conical tube, and 30 μl of 20 mg/ml Proteinase K (Qiagen) were added to the tissue/cell sample and incubated at 55°C overnight. The next day, 30 μl of 10 mg/ml RNAse A (Qiagen) was added to the lysed sample, which was then inverted 25 times and incubated at 37°C for 30 min. Samples were cooled on ice before addition of 2 ml of pre-chilled 7.5 M ammonium acetate (Sigma) to precipitate proteins. The samples were vortexed at high speed for 20 seconds and then centrifuged at ≥ 4,000 g for 10 min. Then, a tight pellet was visible in each tube and the supernatant was carefully transferred into a new 15 ml conical tube. Then, 2 ml chloroform (Sigma) were added and mixed throuhgly well by vortexing. After centrifugation at ≥ 4,000 g for 15 min, the aqueous layer was transferred into a new tube, and then 1 volume of 100% isopropanol (Sigma) was added, inverted 50 times and centrifuged at ≥ 4,000 g for 10 min to pallet DNA. The supernatant was discarded, 6 ml of freshly prepared 70% ethanol was added, the tube was inverted 10 times, and then centrifuged at ≥ 4,000 g for 10 min. The supernatant was discarded by pouring; the tube was briefly spun, and remaining ethanol was removed using a P200 pipette. After air-drying, the DNA changed appearance from a milky white pellet to slightly translucent. Then, 500 μl of ddH2O was added, the tube was incubated at 65°C for 1 h and at room temperature overnight to fully resuspend the DNA. The next day, the gDNA samples were vortexed briefly. The gDNA concentration was measured using a Nanodrop (Thermo Scientific).

### Sequencing readout of APCM sgRNA library representation

Library transduced cells were subjected to genomic DNA (gDNA) extraction using standard molecular biology protocols. The sgRNA library readout was performed using a two-step PCR strategy, where the first PCR includes enough genomic DNA to preserve full library complexity and the second PCR adds appropriate sequencing adapters to the products from the first PCR.

PCR#1 Fwd primer: AATGGACTATCATATGCTTACCGTAACTTGAAAGTATTTCG PCR#1 Rev primer: CTTTAGTTTGTATGTCTGTTGCTATTATGTCTACTATTCTTTCCC

PCR was performed using Phusion Flash High Fidelity Master Mix (PF) (ThermoFisher). In the first round of PCR, we used 3 μg of total gDNA as template input, with thermocycling parameters: 98 °C for 1 min, 16 cycles of (98 °C for 1s, 62 °C for 5s, 72 °C for 30 s), and 72 °C for 2 min. A total of 2 to 4 of PCR#1 reactions was used to capture the full representation of the library in the cells. PCR#1 products for each biological sample were pooled and used for amplification with barcoded second PCR primers. In PCR#2, the thermocycling parameters were: 98 °C for 1 min, 16-18 cycles of (98 °C for 1s, 60 °C for 5s, 72 °C for 30s), and 72 °C for 2 min. Second PCR products were pooled and then normalized for each biological sample before combining uniquely barcoded separate biological samples. The pooled product was then gel purified from a 2% E-gel EX (Life Technologies) using the QIAquick Gel Extraction Kit (Qiagen). The purified pooled library was then quantified with a gel-based method using the Low-Range Quantitative Ladder Life Technologies, dsDNA High-Sensitivity Qubit (Life Technologies), BioAnalyzer (Agilent) and/or qPCR. Diluted libraries with 5-20% PhiX were sequenced with Illumina sequencers.

### Analysis of APCM sgRNA library representation

Raw single-end fastq read files were filtered and demultiplexed using Cutadapt (Martin, 2011). To remove sgRNA scaffold sequences downstream (i.e. 3’ end) of the sgRNA spacer sequences, we used the following command: cutadapt --discard-untrimmed -a GTTTTAGAGCTAGGCCAAC. As the forward PCR primers used to readout sgRNA representation were designed to have a variety of barcodes to facilitate multiplexed sequencing, we then demultiplexed these filtered reads with the following settings: cutadapt -g file:fbc.fasta --no-trim, where fbc.fasta contained the possible barcode sequences within the forward primers. Finally, to remove non-sgRNA sequences upstream (i.e. 5’ end) of the sgRNA spacers, we used the following command: cutadapt --discard-untrimmed –g GTGGAAAGGACGAAACACCG. Through this procedure, the raw fastq read files were trimmed to the 20 bp sgRNA spacer sequences. The 20 bp sgRNA spacer sequences from each demulitplexed sample were mapped the sgRNA spacers to the mSAM library using Bowtie 1.1.2 (Langmead et al., 2009)3/14/23 6:17:00 PM: bowtie -v 2 --suppress 4,5,6,7 --chunkmbs 2000 –best. Using the resultant mapping output, we quantitated the number of reads that had mapped to each sgRNA within the library. The resulting read counts were depth-normalized to log2 reads per million (RPM) in each sample. The sgRNA level abundances were then aggregated to the gene level in each sample. Gene level abundances were then compared across *in vivo* lung metastatic tumors and *in vitro* pre-injection cell pools by linear regression, with 95% confidence intervals shown. Data were represented as mean ± s.e.m, calculated across the experimental replicates within each sample type.

### Tumorigenesis of E0771-APCM in C57BL/6J mice

To generate metastatic lung tumors and perform in vivo screening under immune pressure, 2 × 10^6^ of E0771-mAPCM or E0771-Vector cells were injected into 5-8 weeks old female C57BL/6J mice intravenously. The metastases formation was monitored by IVIS. To generate orthotopic breast tumors, 2 × 10^6^ of E0771 or E0771-dCas9 cells were injected into the orthotopic mammary fat pad of syngeneic 5-8 weeks old female C57BL/6J mice. Tumor sizes were measured every 3-4 days by caliper on the three diameters, and sizes were calculated as approximate spheroids with the formula: Vol = υ/6 *x*y*z. Statistical significance of all tumor growth curves in the study was assessed by analysis of variance (2-way ANOVA), jointly considering the effect of treatment and the passage of time on tumor growth.

### RNA extraction, reverse transcription, and quantitative PCR

RNA in cells were extracted using TRIzol Reagent (Invitrogen) by following standard RNA extraction protocols. The first-strand cDNA of RNA was synthesized using SuperScript™ IV Reverse Transcriptase (Invitrogen). After normalizing the concentrations of cDNA with nuclease-free water, quantitative PCR (qPCR) was performed by adding designated Taqman probe of interested genes, and GAPDH was used as an internal positive control.

### Generation of AAV-ORFs and AAV purification

The polycistronic ORFs of desired genes were cloned into AAV cis plasmid by linearization with AgeI and EcoRI digestion and Gibson assembly. AAVs were produced by co-transfecting HEK293FT cells with above AAV cis plasmids, together with helper plasmid PDF6 and AAV9 or AAV-DJ serotype plasmid to generate AAV-ORFs. Briefly, 5.2 μg of AAV-cis plasmid, 8.7 μg of plasmid AAV9 or AAV-DJ serotype, and 10.4 μg of pDF6 were mixed with PEI at the ratio of 1:3, stand at room temperature for 10-15 min before adding into HEK293FT cells in 150mm-dish at 80-90% confluency. To purify AAV, the transfected HEK293FT cells were collect at days, followed by the addition of 1/10 volume of chloroform, and shake vigorously at 37°C for 1 h. Then, NaCl was added to a final concentration of 1 M and the solution was centrifuged at 20,000 g at 4°C for 15 min. The aqueous layer was collected to another tube, and added PEG8000 to 10% (w/v) and shaken until dissolved. The viral mixture was incubated on ice for 1 h to overnight, and then spun at 20,000 g at 4°C for 15 min. The pellet was resuspended in DPBS and treated with universal nuclease and MgCl2 at 37°C for 30 min. Chloroform (1:1 volume) was added and mixed well by vigorous shaking, and then centrifuged at 12,000 g at 4°C for 15 min. The aqueous layer was collected and concentrated using 100 kDa MWCO ultracentrifuge tubes (Millipore). The genomic copy number (GC) of AAV was determined by real-time quantitative PCR using Taqman probe targeted to the EF1a-Core promoter.

### Therapeutic testing of AAV-ORFs in syngeneic tumor models

Orthotopic breast tumors were established by transplanting 2 × 10^6^ E0771 or E0771-dCas9 cells as specified into the mammary fat pad of 5-8 weeks old female C57BL/6J mice. After transplantation, 1-2×
10^11^ GCs of PBS, AAV-Vector, or AAV-ORF were intratumorally injected to the tumor-bearing mice. Tumors were measured every 3-4 days and tumor sizes were calculated with the formula: Vol = x*y*z*υ/6. Two-way ANOVA and multiple t test were used to compare the size of tumors in treatment groups.

### AAV CAR-T combined treatment in human tumor model in NGS mice

NSG mice were fat pad orthotopically injected with 4-5xe^6^ MCF7 cells with 25% Matrigel at both left and right sites. Mice were intratumorally injected with AAV-Vector at the left site or AAV-MultiImmG at the right site at indicated days. Then 5xe^6^ per dose CAR-T cells were intravenously injected at indicate days.

### Isolation of splenocytes and tumor infiltrating lymphocytes

Syngeneic breast tumors were established by orthotopically transplanting 2 × 10^6^ E0771 cells into the mammary fat pad of 5-8 weeks old female C57BL/6J mice. Tumor-bearing mice were randomly assembled into different treatment groups to receive the treatments as described in the legends. Mice were euthanized at day-30 or other time point as indicated, and the tumors were collected and kept in ice-cold 2% FBS. Splenocytes were washed once with 2% FBS. Tumors were minced into 1-3 mm size pieces using a scalper and then digested using 100 U/mL Collagenase IV for 30-60 min while stirred at 37°C. Tumor suspensions were filtered twice through 100 µm cell strainer, and once through 40 µm cell strainer to remove large bulk masses. RBCs were lysed with 1ml of ACK Lysis Buffer (Lonza) per spleen by incubating 2-5 mins at room temperature, which was followed by the dilution with 10 ml 2% FBS and pass through a 40 µm filter. Splenocytes were resuspended in 2% FBS buffer, counted for flow cytometry staining or RNA isolation. Single cell suspensions of tumors were used for flow cytometry staining, further FACS sorting, or Ficoll purification to obtain tumor-infiltrating lymphocytes.

### Flow cytometry

All antibodies for flow were purchased from Biolegend or eBiosciences. Single-cell suspension of tumor or spleen was prepared using a gentleMACS tissue dissociation system. The panels of antibodies used in the flow cytometry staining as follows: Panel 1: anti-CD45.2-PerCP/Cy5.5, anti-CD3-PE, anti-CD4-PE/Cy7, anti-CD8a-APC/Cy7, anti-PD1-APC or anti-Foxp3-APC, anti-IFN-γ-BV421; Panel 2: anti-CD45-BV421, anti-I-A/I-K-PerCP/Cy5.5, anti-CD11b-FITC, anti-Ly6c-APC; anti-F4/80-PE, anti-CD24-PE/Cy7, anti-PD1-APC/Cy7 anti-CD11c-PE/Dazzle594. All flow antibodies were used at 1:100 dilutions for staining unless otherwise noted. For surface staining, cells were blocked with anti-Fc receptor anti-CD16/CD32, and then stained with surface marker antibodies in the staining buffer of 2% FBS in PBS on ice for 30 min. Samples were washed twice with 2% FBS in PBS before analysis. For intracellular staining, eBioscience™ Intracellular Fixation & Permeabilization Buffer Set was used to fix and permeabilize cells by following manufacturer’s instructions. Briefly, after the staining of surface makers, cells were resuspended in 100 μl Fixation/Permeabilization working solution, and incubated on ice for 10 min before washing with 1× Permeabilization buffer by centrifugation at 600 g for 5 min. Then cell pellet was resuspended in 50 μl 1× Permeabilization buffer with anti-Fc receptor anti-CD16/CD32, and incubated on ice for 10 min, before adding 50 μl 2× intracellular staining antibodies and incubated on ice for 30 min. After staining, cells were centrifuged at 600 g for 5 min, and washed twice with staining buffer before being analyzed or sorted on a BD FACSAria. The data was analyzed using the FlowJo software (v9.9.4 or v10.3). A previously reported strategy was used to define the populations of monocytes, neutrophils, dendritic cell, and macrophages in tumor.

### Sample size determination

Sample size was determined according to the lab’s prior work or similar approaches in the field.

### Randomization and blinding statements

In animal experiments, mice were randomized by sex, cage and littermates. *In vitro* experiments were not randomized. Investigators were blinded in mouse experiments by labeling cages with generic identifiers. Investigators were not blinded d in *in vitro* experiments. In NGS data analysis, investigators were blinded for initial processing of the original data using key-coded metadata.

### Standard statistical analysis

Standard non-NGS statistical analyses were performed in GraphPad Prism using specific statistical tests where appropriate, as detailed in figure legends. NGS statistical analyses were performed in R/RStudio. Different levels of statistical significance were accessed based on specific *p* values and type I error cutoffs (0.05, 0.01, 0.001).

## Supplementary Figure Legends

**Figure S1:**
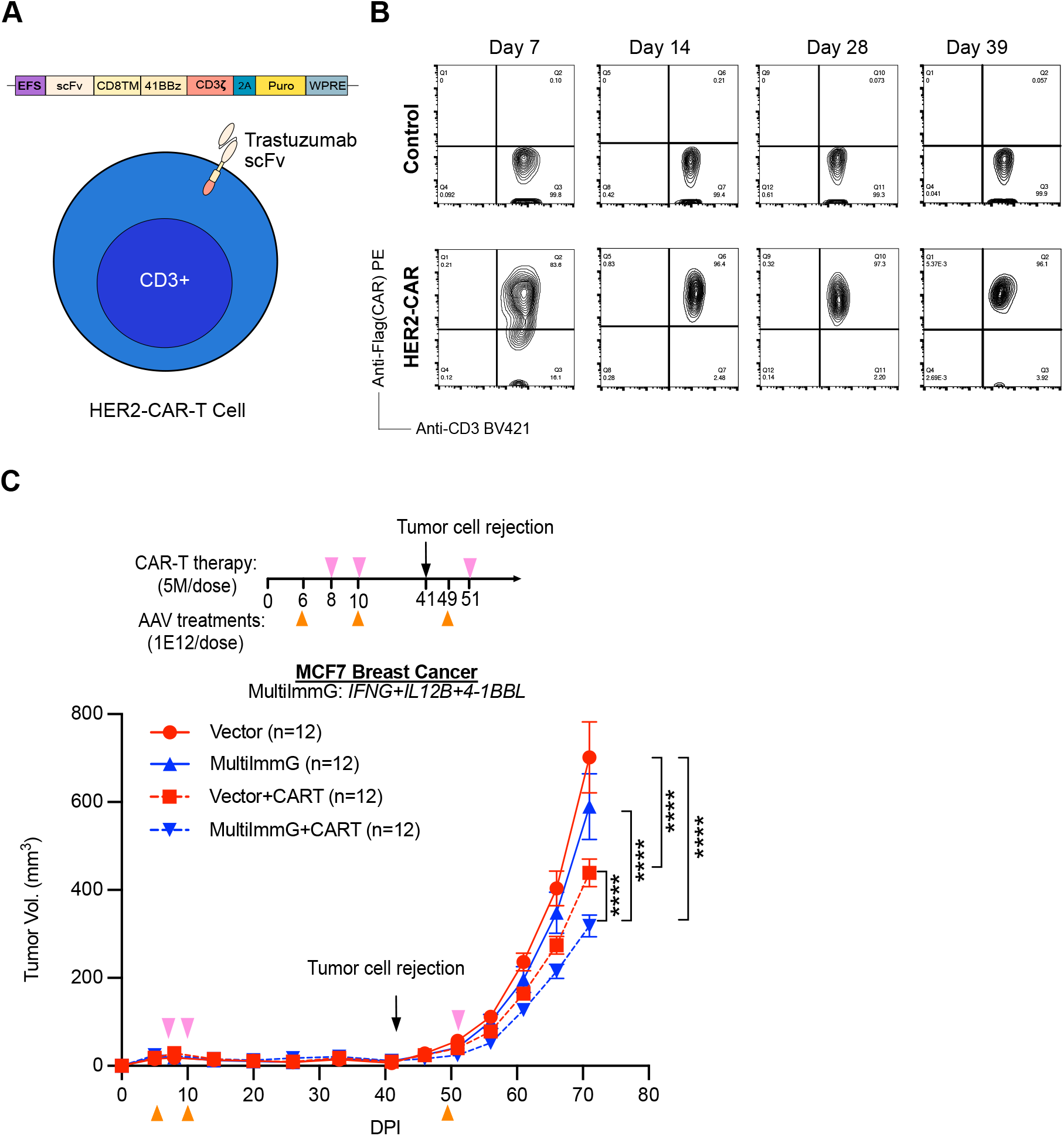
Combination therapy with CAR-T cells and AAV-MultiImmG treatment in solid tumors. **(A)**Schematic diagram for generating HER2-targeting CAR-T cells using the single-chain variant fragment (scFv) from trastuzumab (anti-HER2). CAR-T cells were made by lentiviral infection of peripheral blood mononuclear cells (PBMCs). **(B)**Representative flow gating of CAR-expressing PBMCs 7, 14, 28, and 39 days after lentivirus infection. (C)*In vivo* efficacy of HER2 CAR-T cells and an AAV-MultiImmG combination (IFNG+IL12B+4-1BBL) in a dual-tumor MCF7 human breast cancer rechallenge model. Mice were first intratumorally injected with AAV-Vector at the left site and AAV-MultiImmG at the right site on the indicated days. CAR-T cells were also intravenously injected on the indicated days. Initial tumor were under control for all groups. Mice were rechallenged with MCF7 cells at 41dpi. Data points in this figure are presented as mean ± s.e.m. Statistical significance was assessed by two-way ANOVA. * *p* < 0.05, ** *p* < 0.01, *** *p* < 0.001, **** p < 0.0001. Non-significant comparisons not shown. **Related to: Figure 5**

## Notes

### Competing Interest Statement

The authors have declared no competing interest.

